# Osteoblast-derived Nerve Growth Factor is Required for Skeletal Adaptation to Mechanical Load and the Osteoanabolic Effect of Gambogic Amide in Mice

**DOI:** 10.1101/2025.07.08.663521

**Authors:** Ibtesam Rajpar, Eric McLaughlin, Gabriella Fioravanti, Nicholas Ruggiero, Nohael Cherian, Liliana Minichiello, Ryan E. Tomlinson

## Abstract

In adult mice, new bone accrual following mechanical load is mediated by the neurotrophin nerve growth factor (NGF) that is expressed by osteoblasts on the bone surface. NGF can bind to its high affinity receptor, neurotrophic tyrosine kinase receptor type 1 (TrkA), on peripheral sensory nerves resident in bone and support new bone formation. However, the osteoanabolic therapeutic potential of NGF-TrkA signaling to repair bone is limited due to the long-lasting thermal and mechanical hyperalgesia induced by administration of NGF in mice and humans. Here, we investigated whether 1) mature osteoblasts are the primary source of NGF required for bone accrual following loading, and 2) a small molecule TrkA receptor agonist - gambogic amide - can harness the downstream osteoanabolic potential of NGF-TrkA signaling in the absence of endogenous NGF. Loss of *Ngf* transcription in mature osteoblasts did not appear to affect bone structure or bone mass in adulthood. However, *Ngf* knockout mice significantly reduced periosteal bone accrual and osteogenic *Wnt* transcription in response to loading compared to wildtype mice. Intraperitoneal injection of gambogic amide prior to loading was unable to produce its osteoanabolic effects in *Ngf* knockout mice, suggesting that gambogic amide primarily functions in collaboration with endogenous NGF in bone. In total, our study reveals an important role for osteoblastic NGF in the skeletal adaptation of bone to mechanical forces.

## INTRODUCTION

The skeleton is sensitive and responsive to its mechanical environment (1-3). Indeed, bone cells can convert mechanical cues into biochemical signals in order to direct both anabolic and catabolic processes through the tissue. Generally, in response to mechanical cues, new bone is formed at sites of high strain and removed in areas of low strain. This process of strain adaptive bone remodeling enables the skeleton to alter its bone mass and geometry to meet its functional demands. Furthermore, strain adaptive bone remodeling is an efficient mechanism to maintain the skeleton, by generating bone where it is needed and eliminating bone that is underutilized. Understanding the mechanisms underpinning this well-conserved system is critical for addressing diseases of low bone mass as well as preventing skeletal injuries during extended periods of microgravity or disuse.

One of the best-characterized processes through which tissues perceive and respond to mechanical stimuli is the somatosensory system of peripheral nerves (4), which are distributed throughout the body to transduce signals involved in proprioception and nociception (5). Indeed, primary afferent sensory and sympathetic nerves blanket the surfaces of bone in a dense mesh-like network (6-8). Importantly, the vast majority (> 80%) of nerves in mature bone are thinly myelinated or unmyelinated sensory nerves that express neurotrophic tyrosine kinase receptor 1 (TrkA), the high affinity receptor for nerve growth factor (NGF) (8-11). Thus, the nociceptive nerve fibers in bone are primed to respond to NGF and remain intact in the periosteum throughout life, even as bone undergoes changes associated with aging (12).

In previous studies, we have investigated the role and requirement of NGF-TrkA signaling for strain adaptive bone remodeling (13) as well as the upstream activation of NGF expression in osteoblasts (14). In brief, we reported that osteocalcin-expressing osteoblasts on the surfaces of bone respond to axial forelimb compression with robust NGF expression. Furthermore, inhibition of TrkA signaling significantly decreased load-induced bone formation, whereas administration of exogenous NGF significantly increased load-induced bone formation. However, in addition to causing unwanted side effects – primarily pain (15-20) – NGF is an unlikely candidate for fracture prophylaxis due to the inherent drawbacks of using polypeptides as drugs, including the high cost of production and poor stability.

Therefore, we investigated the use of the small molecule (627.8 g/mol) TrkA agonist gambogic amide (GA) to augment load-induced bone formation. GA was uncovered in a cell-based chemical genetic screen to selectively bind to TrkA and induce tyrosine phosphorylation and downstream signaling (21). Subsequent investigation found that GA induces a lower magnitude but longer lasting activation of TrkA signaling than NGF alone (22). In our previous work, we found that a single dose of GA (0.4 mg/kg) enhanced load-induced bone formation, increased osteogenic gene transcription, and stimulated osteoblast proliferation without obvious pain, as indexed by mechanical and thermal hyperalgesia (23). Importantly, we observed that NGF expression in loaded bone as well as in osteoblastic cells that do not express TrkA was significantly upregulated by GA administration.

Therefore, this study was designed to examine the specific role of osteoblast-derived NGF in load-induced bone formation and response to gambogic amide. Our overall hypothesis was that osteoblasts are the main source of NGF in response to non-damaging mechanical loading and mediate the bulk of the effect of GA in this scenario. To test this hypothesis, we generated a novel mouse model that lacks NGF expression in mature osteoblasts. Next, we assessed the skeleton using microCT and three-point bending, then subjected conditional knockout and wildtype control mice to axial forelimb compression designed to induce lamellar bone formation. Finally, we administered gambogic amide to these mice and assessed its effect in both genotypes. The results from our study reveal how osteoblast-derived NGF mediates the response to mechanical loading and the effect of gambogic amide in the skeleton.

## METHODS

### Mice

Ngf^fl/fl^; Osteocalcin (OC)-Cre+ male and female mice (cKO) on a C57BL/6J background were generated for this study in accordance with the IACUC of Thomas Jefferson University. Mice with homozygous floxed *Ngf* alleles (24) were crossed with mice carrying Cre recombinase driven by the osteocalcin promoter (Jackson Labs #019509). Wildtype (Ngf^fl/fl^) littermates were included as controls in the study. All mice were housed at room temperature and fed a daily diet of rodent chow (LabDiet 5001).

### *In vivo* mechanical loading

Axial forelimb compression of the right forelimb was performed using a material testing system (TA Instrument Electroforce 3200 Series III) with custom designed fixtures. Prior to loading, mice were anesthetized with 3% isoflurane and buprenorphine (0.12mg/kg, IP), and they were maintained under 1.5% isoflurane for the duration of the loading protocol. Each mouse was loaded using a 2Hz sinusoidal, rest-inserted waveform with a peak force of 3N for 100 cycles each day over three consecutive days (D0-D2). This loading protocol has previously been shown to induce robust lamellar bone formation at the mid-shaft of the ulna (23, 25). The non-loaded left forelimb served as a contralateral control. In a separate group of WT and cKO mice, gambogic amide (Enzo #BML-N159) was administered via IP injection (0.4 mg/kg in 100 ul of 10% DMSO) one hour prior to loading.

### Histomorphometry

To assess load-induced bone formation by dynamic histomorphometry, adult (16-20 weeks) male and female mice were given calcein (10mg/kg, Sigma C0875) on D3 and alizarin red s (30mg/kg, Sigma A3882) on D8 of the loading protocol by IP injection. Animals were euthanized on D10. Forelimbs were dissected, fixed in 10% formalin overnight at 4°C, dehydrated in 70% ethanol, and embedded in polymethylmethacrylate. 100 micron thick sections from the mid-diaphysis of the ulnae were cut using a precision saw (Isomet 1000) and mounted on glass slides with Eukitt mounting medium (Sigma 03989) and visualized by confocal microscopy (Zeiss). Bone formation parameters, including endosteal (Es) and periosteal (Ps) mineralizing surface (MS/BS), mineral apposition rate (MAR), and bone formation rate (BFR/BS), were quantified using a custom ImageJ plugin according to the guidelines of the ASBMR Committee for Histomorphometry Nomenclature (26).

### microCT imaging and three-point bending

For analysis of skeletal phenotype by microCT, the right femurs were dissected from adult mice and frozen at - 20°C in PBS-soaked gauze. Each bone was scanned using a Bruker microCT analyzer fixed with a 1 mm aluminum filter, and with scanning parameters of 55 kV and 181 uA at a resolution of 12 microns. Bone scans were reconstructed using nRecon (Bruker), and aligned and analyzed using CTan (Bruker) for cortical and trabecular bone parameters. For mechanical testing of adult femurs with three-point bending, femurs were placed on custom fixtures with the condyles facing down and a span length of 7.6 mm measured with calipers. Femurs were then subjected to a monotonic displacement ramp of 0.1 mm/s until failure. Force and displacement data from the test as well as geometric parameters from microCT were analyzed using a custom GNU Octave script to derive structural and material properties.

### RNA isolation and gene expression analysis

The middle third of ulnae from adult, male mice were harvested 3 and 24 hours after a single bout of compressive loading on D0 to extract RNA for gene expression analysis. Bones were centrifuged to remove bone marrow, then stored in RNAlater (ThermoFisher) at -80 °C. Bone tissue was pulverized in TRIzol (ThermoFisher) to obtain uniform tissue homogenates, and total RNA was isolated from bone homogenates using the TRIzol method as per manufacturer’s instructions. RNA was further purified using a Qiagen RNeasy kit (Qiagen), and quantified using a NanoDrop spectrophotometer. RNA was reverse transcribed using iScript cDNA synthesis kit (BioRad), and amplified using SYBR Green biochemistry with the QuantStudio 3 real time PCR system (Applied Biosystems). Relative gene expression was quantified using the ΔΔCT method, using GAPDH as a reference gene. Primer sequences were designed using Primer-BLAST and purchased from IDT Technologies. A complete list of primer sequences is provided in **Table 1**.

**Table 1.**
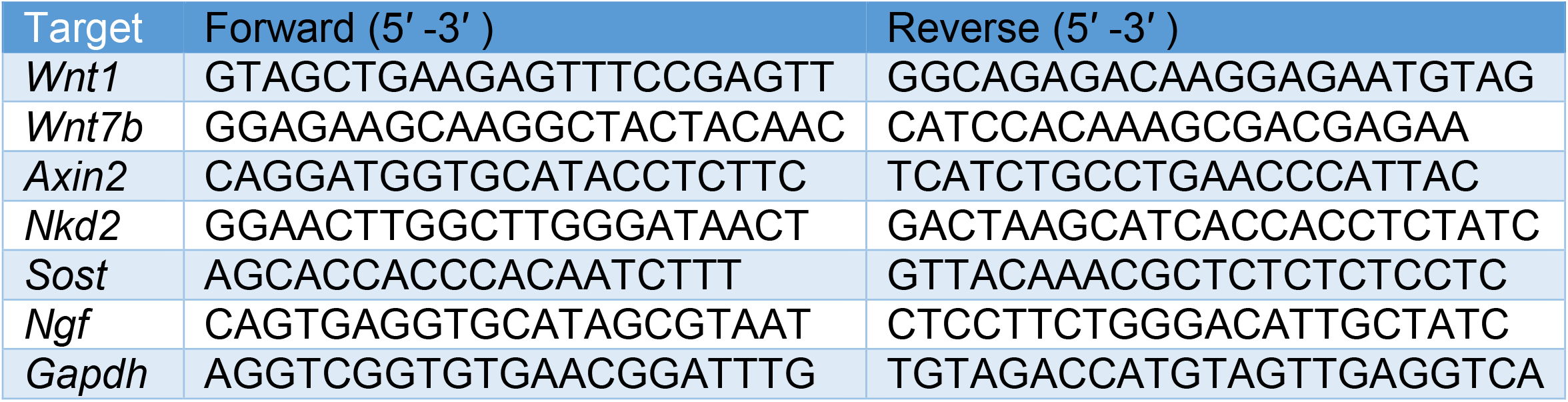
Oligonucleotide primers used for qRT-PCR.

### Statistical analysis

Data was collected in a blinded fashion from all animals in the study, unless removed prior to analysis due to illness or injury after consultation with veterinarian staff. After data collection, outlier analysis was performed using Grubbs’ test if necessary. For *in vivo* comparisons of bone formation parameters in loaded versus non-loaded bone, two-way ANOVA was used with post hoc Tukey’s tests. For analysis of skeletal phenotype, two-tailed, unpaired t-tests were used (adjusted p ≤ 0.05). Statistical analysis was carried out in Prism 10 (Graphpad).

## RESULTS

### NGF conditional knockout mice have normal adult skeleton

To determine the role of *Ngf* in skeletal development and adult bone integrity, we generated conditional knockout mice in which *Ngf* was selectively deleted from the osteoblast lineage using Osteocalcin-Cre. Microcomputed tomography (microCT) analysis of femoral bones from adult conditional knockout (cKO) and littermate control mice revealed no significant differences in tibial cortical bone (**Fig. 1**) or vertebral trabecular bone (**Fig. 2**) architecture, including parameters such as bone area, cortical thickness, or bone volume fraction. To assess bone mechanical properties, we performed three-point bending (3PB) tests on femoral midshafts. No differences were observed in stiffness, ultimate force, or work to failure between Ngf cKO and control bones (**Fig. 3**). Together, these findings indicate that loss of *Ngf* transcription in osteoblast-lineage cells during development does not impair skeletal morphology, mass, or mechanical strength in adulthood.

**Figure 1.**
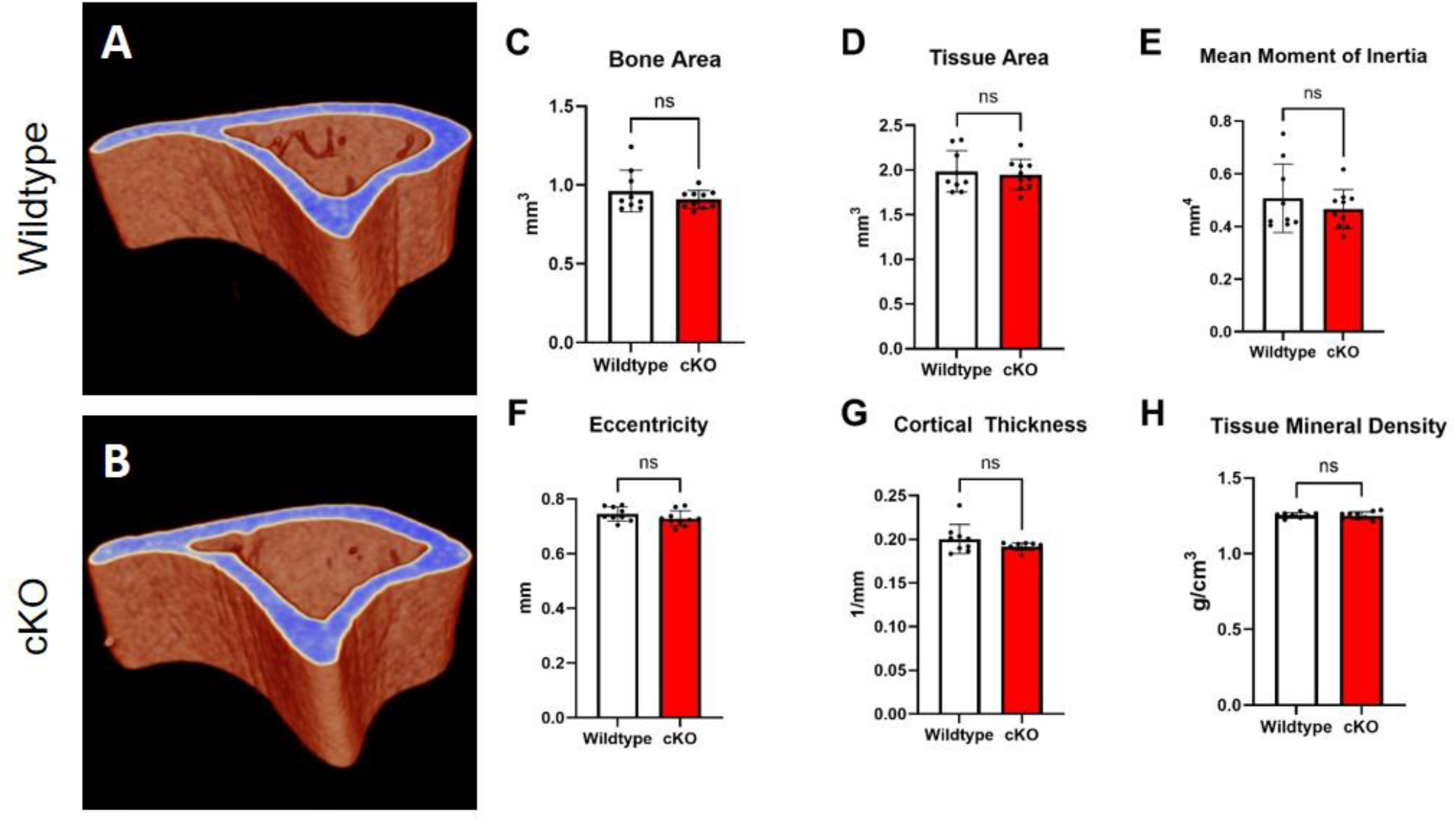
MicroCT analysis of tibial midshaft. Analysis of 1 mm of tibial cortical bone by microCT from A) wildtype and B) cKO mice revealed no differences between genotypes in C-H) cortical parameters.

**Figure 2.**
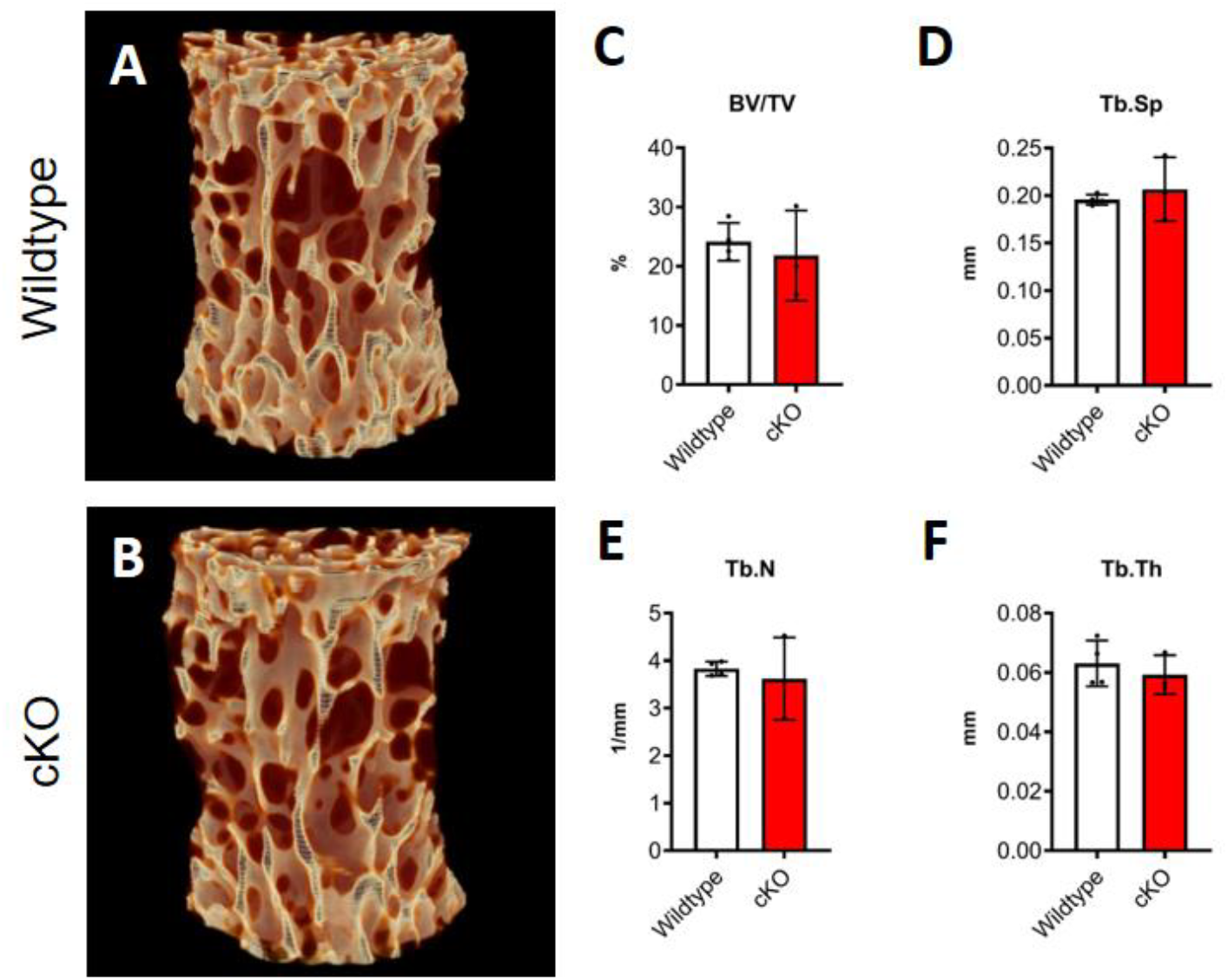
MicroCT analysis of lumbar spine. Analysis of entire L5 vertebra by microCT from A) wildtype and B) cKO mice revealed no differences between genotypes in C-F) trabecular bone parameters.

**Figure 3.**
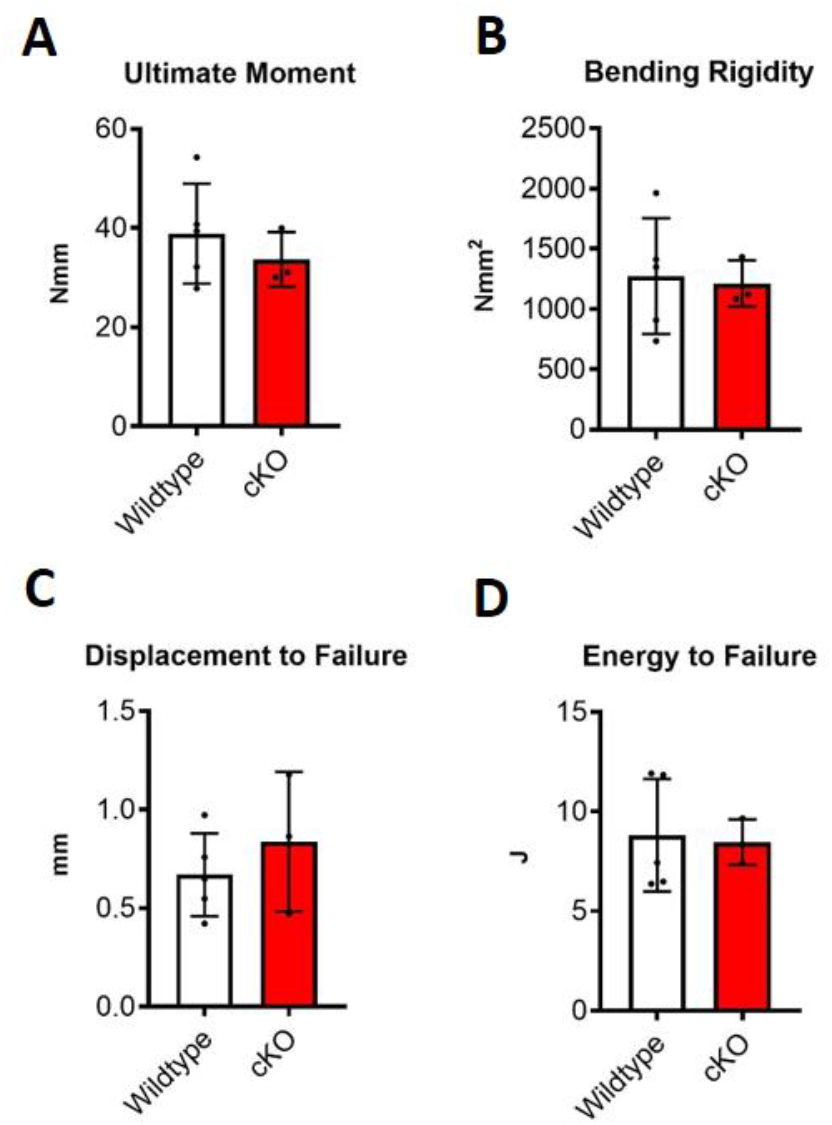
Three-point bending of femur. Analysis of mechanical properties of femur by three-point bending revealed no significant differences in A) ultimate moment B) bending rigidity C) displacement to failure or D) energy to failure.

### Osteoblastic NGF is required for normal load-induced bone formation

To assess whether osteoblastic NGF contributes to the skeletal response to mechanical loading, we subjected adult conditional knockout and control mice to a single bout of axial forelimb compression and analyzed bone formation by dynamic histomorphometry. While control mice exhibited a robust anabolic response to loading, characterized by increased mineralizing surface (MS/BS), mineral apposition rate (MAR), and bone formation rate (BFR/BS), NGF cKO mice showed a blunted response (**Fig. 4**). The most prominent deficit in the cKO animals was a significant reduction in MAR, suggesting that osteoblastic *Ngf* is required for translating mechanical stimuli into matrix mineralization rather than influencing the extent of surface activation. These results indicate that *Ngf* transcription in osteoblast-lineage cells is essential for load-induced bone formation.

**Figure 4.**
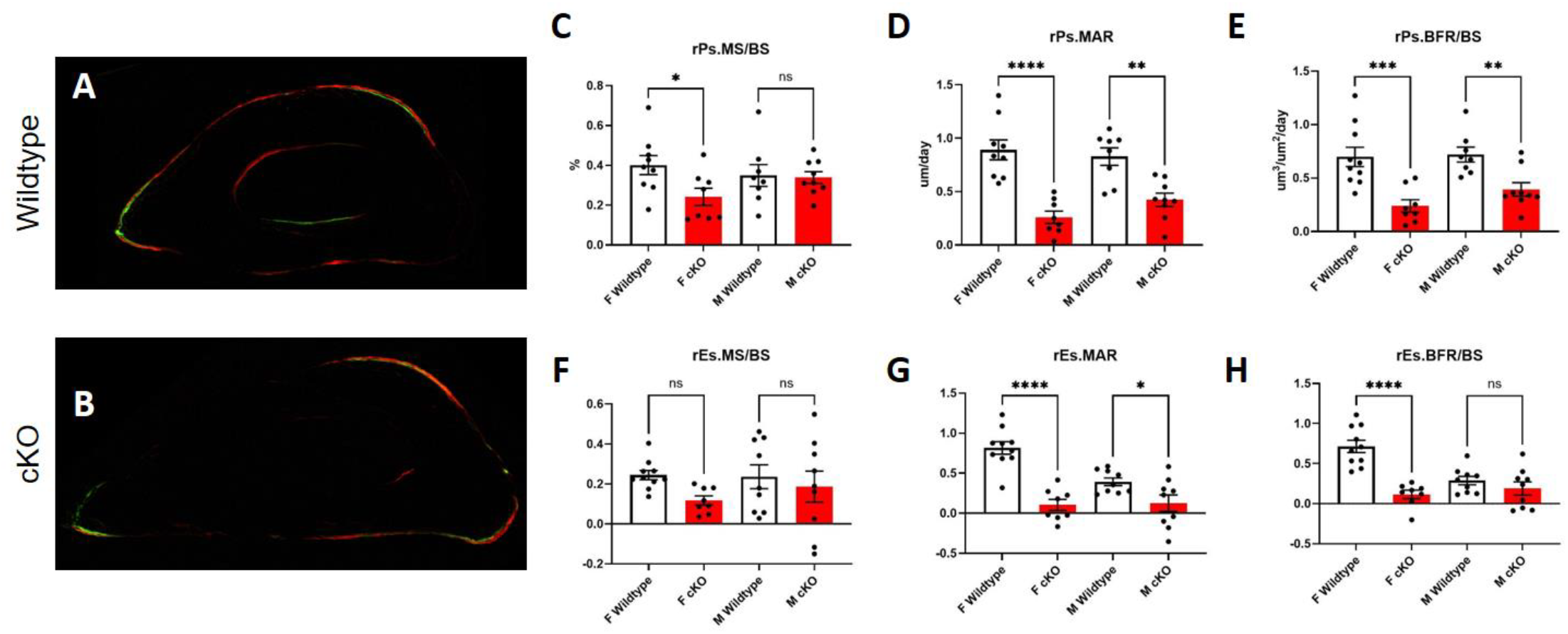
Loss of NGF impairs load-induced bone formation. Dynamic histomorphometry on A) wild type and B) conditional knockout mice was performed following administration of calcein (green) and alizarin red (red) labels, which revealed significant differences between genotypes in relative (loaded – non-loaded) C-D) periosteal and F-H) endosteal bone formation parameters for both male (M) and female (F) mice. * p < 0.05 vs. control.

### Osteoblastic NGF drives osteoanabolic Wnt signaling in the early response to mechanical load

To investigate the molecular basis of this impaired anabolic response, we performed qRT-PCR on cortical bone from loaded and non-loaded limbs at 3 and 24 hours post-compression. In control mice, mechanical loading induced the expected upregulation of *Ngf* at both time points, consistent with its role as a mechanoresponsive gene in osteoblasts (**Fig. 5**). This induction was absent in cKO mice, confirming the specificity of the knockout. Expression of *Wnt7b*, as well as Wnt/β-catenin target genes *Axin2* and *Nkd2*, was significantly diminished in cKO mice at both 3 and 24 hours, indicating impaired activation of osteogenic Wnt signaling in the absence of *Ngf*. While *Sost* levels were unchanged at 3 hours, they were significantly elevated at 24 hours in cKO bone, potentially contributing to suppressed Wnt activity. Notably, *Wnt1*, another mechanoresponsive Wnt ligand, was only reduced at 24 hours in the mutant. These findings support a model in which osteoblastic *Ngf* is necessary for the early induction and maintenance of Wnt signaling in response to mechanical loading, enabling proper bone formation.

**Figure 5.**
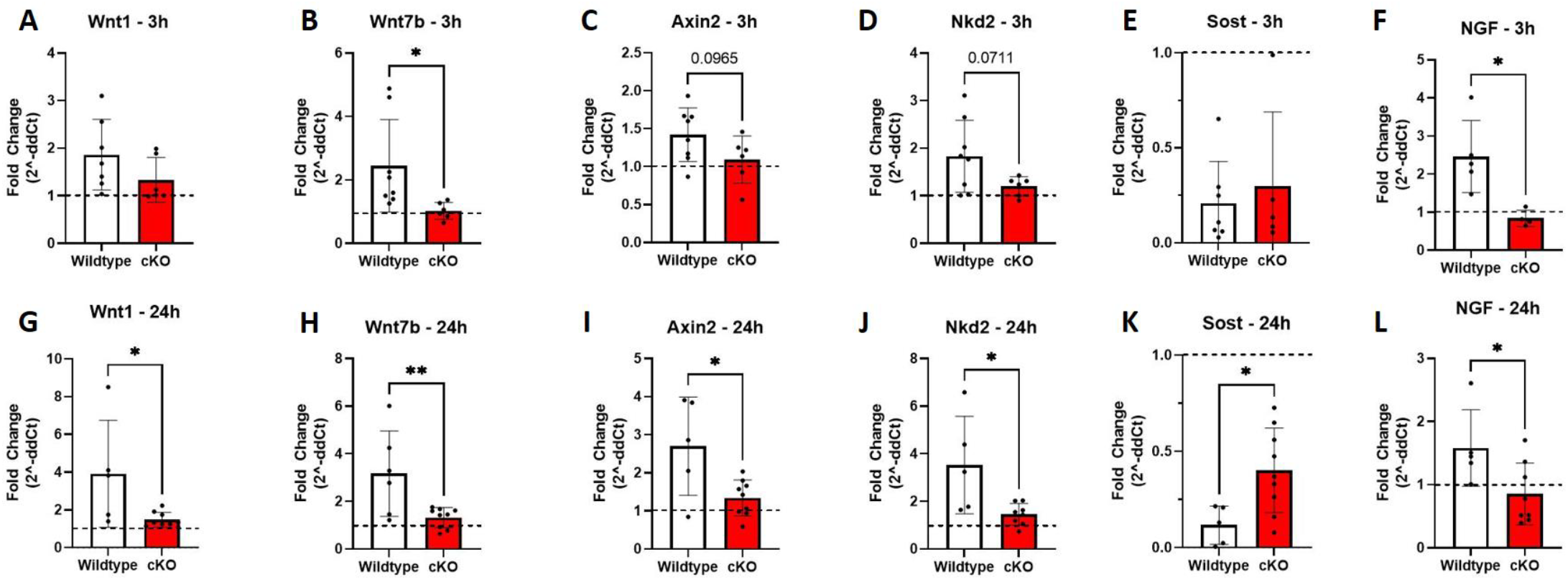
Gene expression in loaded forelimbs. qRT-PCR was used to determine the effect of the loss of NGF in the osteoblast lineage following axial forelimb compression at A-F) 3 hours and G-L) 24 hours after loading. Notably, *Wnt7b* and *Ngf* were significantly diminished at 3 hours, with *Wnt1, Wnt7b, Axin2, Nkd2, Sost*, and *Ngf* significantly different at 24 hours. * p < 0.05 vs wildtype.

### Gambogic Amide Enhances Load-Induced Bone Formation in Wildtype but Not Ngf cKO Mice

To further investigate the role of osteoblastic Ngf in mediating the anabolic effects of NGF-TrkA pathway activation, we conducted an additional experiment using only female mice, as they exhibited the strongest phenotype in the conditional knockout model. Wildtype and Ngf cKO mice were treated with either vehicle or the TrkA agonist gambogic amide (GA) and subjected to axial forelimb compression. Dynamic histomorphometry revealed that GA treatment enhanced bone formation in response to loading in wildtype mice, as evidenced by a significant increase in relative periosteal bone formation rate (rPs.BFR/BS), while no such enhancement was observed in Ngf cKO mice (**Fig. 6**). Although mineral apposition rate (MAR) was the primary driver of overall changes, there were no significant differences in MAR between GA-and vehicle-treated groups within either genotype. A similar trend was observed on the endosteal surface, where GA-treated wildtype mice showed a non-significant increase in BFR/BS (p < 0.10), whereas no effect was detected in the Ngf cKO group. These findings suggest that osteoblast-derived NGF is required for the full anabolic response to GA-mediated TrkA activation during mechanical loading.

**Figure 6.**
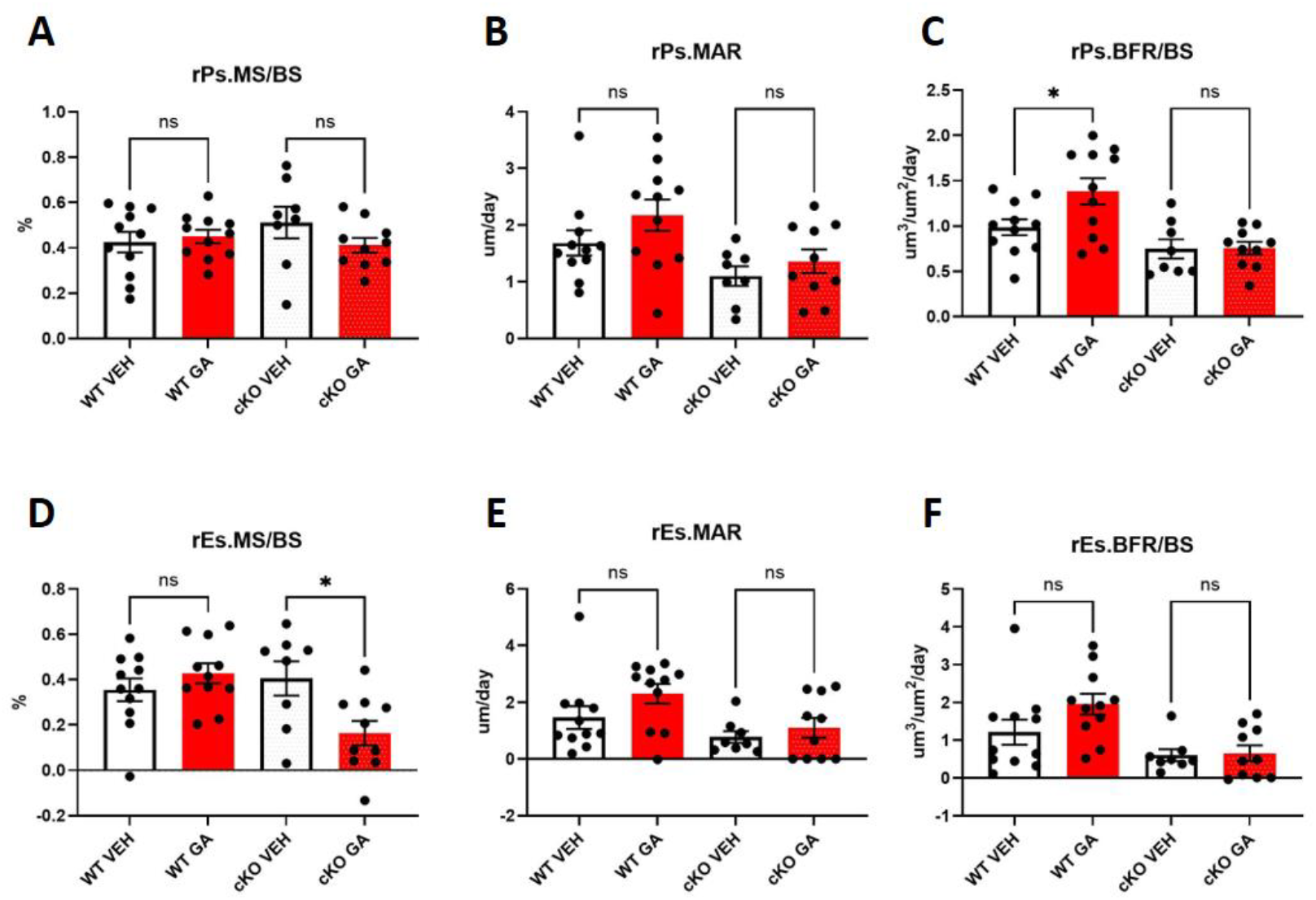
Gambogic amide does not improve load induced bone formation in cKO mice. In female wildtype (WT) and NGF conditional knockout (cKO) mice administered vehicle (VEH) or gambogic amide (GA), relative (loaded – non-loaded) A-C) periosteal and D-F) endosteal bone formation parameters were quantified by dynamic histomorphometry. * p < 0.05 vs vehicle.

## DISCUSSION

In this study, we defined the role of osteoblast-derived nerve growth factor (NGF) in skeletal adaptation using a conditional knockout (cKO) mouse model. Specifically, we found that deletion of *Ngf* from the mature osteoblast lineage did not affect bone mass, architecture, or mechanical properties in adult mice. However, cKO mice exhibited a significantly blunted anabolic response to mechanical loading, demonstrating the key role for osteoblast-derived NGF in load-induced bone formation. Furthermore, we found that administration of gambogic amide (GA), a selective small molecule TrkA agonist, enhanced load-induced bone formation in WT mice but not cKO mice. Together, these findings suggest that osteoblastic NGF is dispensable for normal skeletal maintenance but required for load-induced bone formation and the efficacy of GA on bone.

Our previous work has demonstrated that NGF-TrkA signaling is required to support load-induced bone formation in response to loading through interactions between sensory nerves and bone cells (13). The results from this study extend our model by identifying mature osteoblasts as the primary, if not only, source of NGF in bone following non-damaging osteogenic mechanical loading. Furthermore, our transcriptional profiling revealed that loss of osteoblastic NGF disrupted expression of Wnt ligands (notably *Wnt7b* and *Wnt1*) and target genes (*Axin2, Nkd2*), indicating that osteoblast-derived NGF signaling is upstream of Wnt-mediated osteoanabolic signaling and consistent with previous work inhibiting TrkA signaling directly (13). Future studies may investigate whether osteoblastic NGF is able to influence *Sost* expression, potentially indirectly through activation of TrkA expressing sensory nerves or other cells.

In this study, we also tested whether gambogic amide (GA), a small-molecule TrkA agonist, acts independently of endogenous osteoblastic NGF. In wildtype mice, GA enhanced periosteal bone formation following mechanical loading, consistent with our previous report (23). In contrast, the effect of GA on load-induced bone formation was entirely absent in NGF cKO mice. This result suggests that GA does not activate TrkA signaling in the absence of NGF, but rather augments an NGF-initiated response. One possibility is that GA requires NGF-bound TrkA receptors to exert its effect, perhaps by stabilizing receptor dimerization to prolong signaling duration. Alternatively, we previously reported that GA increases NGF expression in osteoblasts (23), but GA would not be able to exert this effect in cKO mice. Nonetheless, these results suggest that GA does not fully mimic NGF as a TrkA agonist in bone but rather functions in concert with endogenous NGF to support NGF-TrkA signaling.

We acknowledge this study has some limitations. First, this study relies on the use of a single Cre driver (Osteocalcin-Cre or OC-Cre, also known as BGLAP-Cre (27)) to remove *Ngf* from the mature osteoblast lineage. However, OC-Cre recombination has previously been observed in multiple non-skeletal tissues, such as muscle and brain (28), representing a critical caveat regarding our observation that osteoblast-derived NGF is the sole source of NGF in our loading experiments. Future studies using inducible or broader Cre lines may help clarify this result. In addition, we focused on female mice for our GA studies due to the stronger cKO phenotype. However, we acknowledge the possibility that GA may affect male mice differently. Finally, our transcriptional analysis supports impaired Wnt signaling as a key downstream effect of osteoblast-derived *Ngf*, but the direct mechanistic link between load-induced NGF-TrkA signaling and Wnt pathway activation in bone remains to be elucidated.

## CONCLUSION

In summary, our findings demonstrate that NGF produced by mature osteoblasts is essential for normal skeletal adaptation to mechanical loading, acting as an upstream regulator of Wnt signaling and is necessary to support the efficacy of GA in bone. These results clarify the cellular origin and signaling role of NGF in bone mechanotransduction and support further investigation into therapeutic strategies that harness NGF-TrkA signaling without the drawbacks of exogenous NGF administration. Selective activation or augmentation of osteoblastic NGF, or its downstream pathways, may offer new avenues for promoting bone formation in disuse, aging, or microgravity-related bone loss.

## ACKNOWLEDGEMENTS

This research was supported by the National Institute of Arthritis and Musculoskeletal and Skin Diseases under award number AR074953 (RET). The content is solely the responsibility of the authors and does not necessarily represent the official views of the funding bodies. The authors have no conflicts of interest to disclose.

## DECLARATION OF INTERESTS

The authors declare that they have no known competing financial interests or personal relationships that could have appeared to influence the work reported in this paper.

## DATA AVALIBILITY STATEMENT

All data will be made available from the investigators upon request.

## Notes

### Competing Interest Statement

The authors have declared no competing interest.

